# An ERK5-KLF2 signalling module regulates early embryonic gene expression dynamics and stem cell rejuvenation

**DOI:** 10.1101/2021.03.29.435803

**Authors:** Helen A. Brown, Charles A.C. Williams, Houjiang Zhou, Diana Rios-Szwed, Rosalia Fernandez-Alonso, Saria Mansoor, Liam McMulkin, Rachel Toth, Robert Gourlay, Julien Peltier, Nora Dieguez-Martinez, Matthias Trost, Jose M. Lizcano, Marios P. Stavridis, Greg M. Findlay

## Abstract

The ERK5 MAP kinase signalling pathway drives transcription of naïve pluripotency genes in mouse Embryonic Stem Cells (mESCs). However, how ERK5 impacts on other aspects of mESC biology has not been investigated. Here, we employ quantitative proteomic profiling to identify proteins whose expression is regulated by the ERK5 pathway in mESCs. This reveals a function for ERK5 signalling in regulating dynamically expressed early embryonic 2-cell stage (2C) genes including the mESC rejuvenation factor ZSCAN4. ERK5-dependent ZSCAN4 induction in mESCs increases telomere length, a key rejuvenative process required for prolonged culture. Mechanistically, ERK5 promotes ZSCAN4 and 2C gene expression via transcription of the KLF2 pluripotency transcription factor. Surprisingly, ERK5 also directly phosphorylates KLF2 to drive ubiquitin-dependent degradation, encoding negative-feedback regulation of 2C gene expression. In summary, our data identify a regulatory module whereby ERK5 kinase and transcriptional activities bi-directionally control KLF2 levels to pattern dynamic 2C gene transcription and mESC rejuvenation.

## Introduction

Embryonic Stem Cells (ESCs) can self-renew or differentiate along any lineage in the adult body, a property known as pluripotency (Nichols and Smith, 2009). The fundamental regulatory mechanisms which govern pluripotency are therefore of intense interest to exploit pluripotent cells in regenerative therapeutics (Shi et al., 2017). Cellular signalling network activity plays a critical role in ESC decision-making, by implementing specific gene expression signatures to define ESC developmental choice. Although many signalling networks relevant for ESC decision-making have been identified, it remains a challenge to understand the molecular and biological functions of critical ESC signalling pathways in regulating pluripotency and lineage-specific transcriptional networks (Fernandez-Alonso et al., 2017).

Recently, ERK5 (also known as BMK1) (Lee et al., 1995; Zhou et al., 1995) was identified as a key regulator of transition between naïve and primed pluripotency in mouse ESCs (mESCs) (Williams et al., 2016). ERK5 signalling promotes the naïve state by driving expression of pluripotency genes (Williams et al., 2016), although the wider molecular targets and biological functions of the ERK5 pathway in mESCs have not been identified. In this regard, ERK5 uniquely encodes a kinase domain and a putative transcriptional activation domain (Kasler et al., 2000), both of which are required to support pluripotency gene expression (Williams et al., 2016). Therefore, current data suggest that catalytic and transcriptional functions of ERK5 are to likely play a role in mediating ERK5 function in mESCs (Figure 1A). However, ERK5 substrates and wider transcriptional networks have not yet been explored in this context.

**Figure 1.**
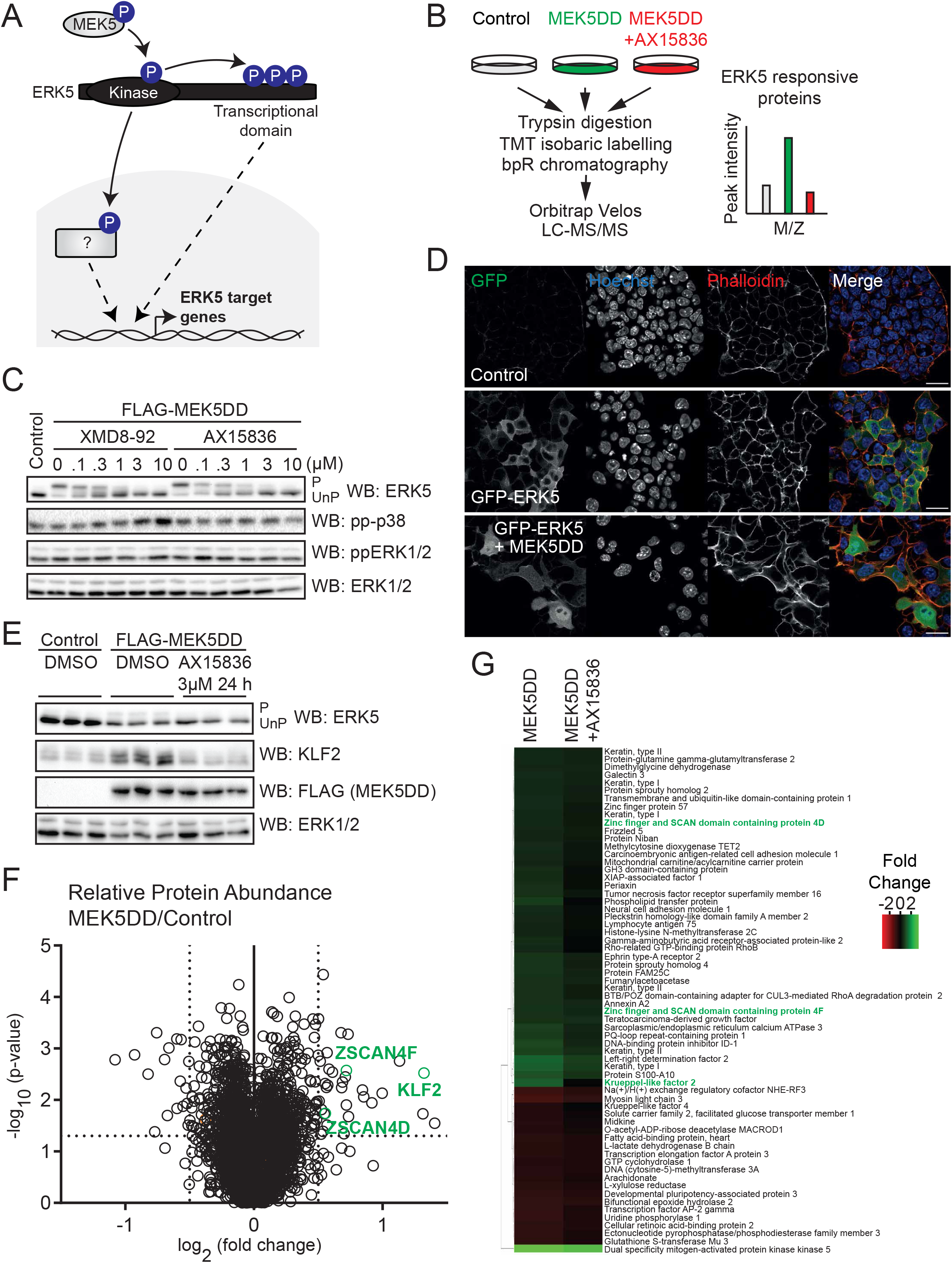
Quantitative analysis of ERK5-dependent proteome dynamics in mESCs. (A) ERK is phosphorylated and activated by MEK5, leading to phosphorylation of transcription factor substrates or autophosphorylation of a putative C-terminal transcription activation domain. These mechanisms together are hypothesised to control ERK5-dependent gene expression and proteome dynamics. (B) Quantitative proteomics workflow: Wildtype mESCs were transfected (48 h) with empty vector (Control), constitutively-active MEK5 (MEK5DD), or MEK5DD and treated the selective ERK5 inhibitor AX15836 (3 μM, 24 h prior to lysis). Cells were lysed, proteins digested with trypsin, TMT labelled and analysed by LC-MS/MS. Proteins responsive to ERK5 signalling are predicted to be induced by MEK5DD, and this effect reversed by AX15836. (C) mESCs were transfected with MEK5DD to (48 h) and treated with indicated concentrations of selective ERK5 inhibitors XMD8-92 and AX15836 (24 h). ERK5 activation was assessed by band-shift following ERK5 immunoblotting (P = phosphorylated activate ERK5, UnP = unphosphorylated inactive ERK5), and p38 MAPK and ERK1/2 activation assessed by immunoblotting with phosphospecific antibodies. ERK1/2 levels are used as a loading control. (D) mESCs were transfected with empty vector (Control), GFP-ERK5 or GFP-ERK5 and MEK5DD (24 h) before fixation. Anti-GFP antibody was used to determine ERK5 localisation. Phalloidin was used to visualise cell boundaries, and Hoechst to stain cell nuclei. Scale bar = 25 μm. (E) mESCs were transfected with either empty vector control or MEK5DD (48 h) and treated with AX15836 (3 μM, 24 h prior to lysis), with parallel samples used for proteomic analysis. ERK5 activation was assessed by band-shift following ERK5 immunoblotting (P = phosphorylated activate ERK5, UnP = unphosphorylated inactive ERK5) and by KLF2 induction. MEK5DD expression was determined by immunoblotting for the FLAG tag. ERK1/2 was used as a loading control. (F) mESCs were transfected with either empty vector control or MEK5DD (48 h) and treated with either DMSO or AX15836 (3 μM, 24 h prior to lysis), and relative protein abundance quantified by mass-spectrometry. Volcano plot shows log_2_ (fold change) of protein abundance against –log_10_ (p value). Thresholds were set as >0.5 log_2_ (fold change), and p value <0.05. KLF2 (green) is shown as a positive control. Expressed constitutively-active MEK5DD is omitted from the volcano plot for clarity. (G) Abundance of proteins above >0.5 log_2_ (fold change) and p value <0.05 thresholds relative to control shown in (F) represented by heat map.

Here, we use state-of-the-art quantitative proteomics to identify the ERK5-responsive proteome in mESCs. Within a specific cohort of ERK5-dependent proteins is ZSCAN4, a key member of a network of early embryonic 2-cell stage specific (2C) genes (Falco et al., 2007) that promotes attainment of naïve pluripotency (Hirata et al., 2012; Zhao et al., 2018) and stem cell ‘rejuvenation’ *in vitro* (Amano et al., 2013). We show that ERK5 induction of the key pluripotency transcription factor KLF2 mediates expression of *Zscan4* and other 2C genes. Furthermore, ERK5 signalling and ZSCAN4 induction promote telomere elongation, a key process that contributes to mESC rejuvenation. Unexpectedly, we find that ERK5 also directly phosphorylates KLF2 at dual pSer/Thr-Pro motifs, which recruits a Cullin family E3 ubiquitin ligase to promote KLF2 ubiquitylation and proteasomal degradation. KLF2 phosphorylation thereby enables a negative feedback loop to suppress expression of *Zscan4* and other 2C genes. In summary, our data provide molecular insight into ERK5 kinase and transcriptional functions in mESCs, which directionally modulate KLF2 levels to pattern early embryonic 2C gene transcription and mESC rejuvenation.

## Results

### Quantitative proteomic profiling identifies ERK5 regulated proteins in mESCs

ERK5 signalling drives transcription of naïve pluripotency genes in mESCs (Figure 1A) (Williams et al., 2016). However, the wider impact of the ERK5 pathway on gene expression and proteome dynamics in mESCs remains uncertain. In order to tackle this question, we set out to systematically identify proteins whose expression is regulated by ERK5 signalling. To this end, we developed a quantitative proteomics workflow employing complementary strategies to specifically activate and inhibit ERK5 signalling in mESCs (Figure 1B). As the ligands that activate ERK5 in mESCs are not yet known, we employed a constitutively active mutant of the specific upstream kinase MEK5 (MEK5DD) (English et al., 1999; Kato et al., 1997) to specifically activate ERK5, and the selective ERK5 inhibitors XMD8-92 (Yang et al., 2010) and AX15836 (Lin et al., 2016) to inhibit ERK5 kinase activity.

As proof-of-principle, we demonstrate highly sensitive manipulation of ERK5 activity using C-terminal autophosphorylation and resulting retarded electrophoretic mobility as a readout (Kato et al., 1997). MEK5DD expression in mESCs activates ERK5, and this is reversed by treatment with the selective ERK5 inhibitors XMD8-92 and AX15836 (Figure 1C). Importantly, the related MAP kinases ERK1/2 and p38 are not significantly activated or inhibited by modulation of the ERK5 pathway (Figure 1C). Furthermore, ERK5 activation is accompanied by translocation to the nucleus (Figure 1D), and expression of the pluripotency transcription factor KLF2, a known ERK5 target gene (Morikawa et al., 2016; Parmar et al., 2006; Sohn et al., 2005; Sunadome et al., 2011; Williams et al., 2016), is dynamically regulated by ERK5 activity (Figure 1E). These data confirm that our experimental approach robustly and specifically modulates ERK5 kinase activity and gene expression in mESCs.

These defined conditions for ERK5 activation and inhibition allowed us to conduct quantitative proteomic profiling to elucidate the ERK5-regulated proteome in mESCs. This experimental approach identified a total of 8732 proteins, of which 7639 were quantified by at least 2 unique peptides. The abundance of 56 proteins changes >0.5 (log_2_) upon ERK5 activation by MEK5DD (Figure 1F, G), indicating that ERK5 signalling acutely and selectively regulates proteome dynamics. Furthermore, ERK5 inhibition by AX15836 largely suppresses proteins that are induced upon ERK5 activation (Figure 1G), indicating a key role for ERK5 kinase activity. KLF2, which is a transcriptional target of ERK5 signalling (Morikawa et al., 2016; Parmar et al., 2006; Sohn et al., 2005; Sunadome et al., 2011; Williams et al., 2016), is significantly induced in response to ERK5 pathway activation as expected (Figure 1F, G). Amongst proteins whose expression is significantly induced by ERK5 signalling are ZSCAN4D and ZSCAN4F (Figure 1F, G), which are expressed in the early embryonic 2-cell stage (2C) and plays a central role in mESC genome stability and rejuvenation (Amano et al., 2013; Falco et al., 2007; Hirata et al., 2012; Zalzman et al., 2010; Zhao et al., 2018).

### The ERK5 pathway drives expression of ZSCAN4 and other 2C-stage genes

As a functional connection between ERK5 signalling and ZSCAN4 expression has not been reported, we first set out to validate the role of ERK5 in regulating ZSCAN4 expression. Analysis of ZSCAN4 protein levels by immunoblotting confirms that ERK5 activation by MEK5DD induces ZSCAN4 expression, and this is reversed by treatment with AX15836 (Figure 2A). This mirrors regulation of known ERK5 pathway target KLF2 (Figure 2A), confirming ZSCAN4 as a novel protein target of the ERK5 signalling pathway in mESCs.

**Figure 2.**
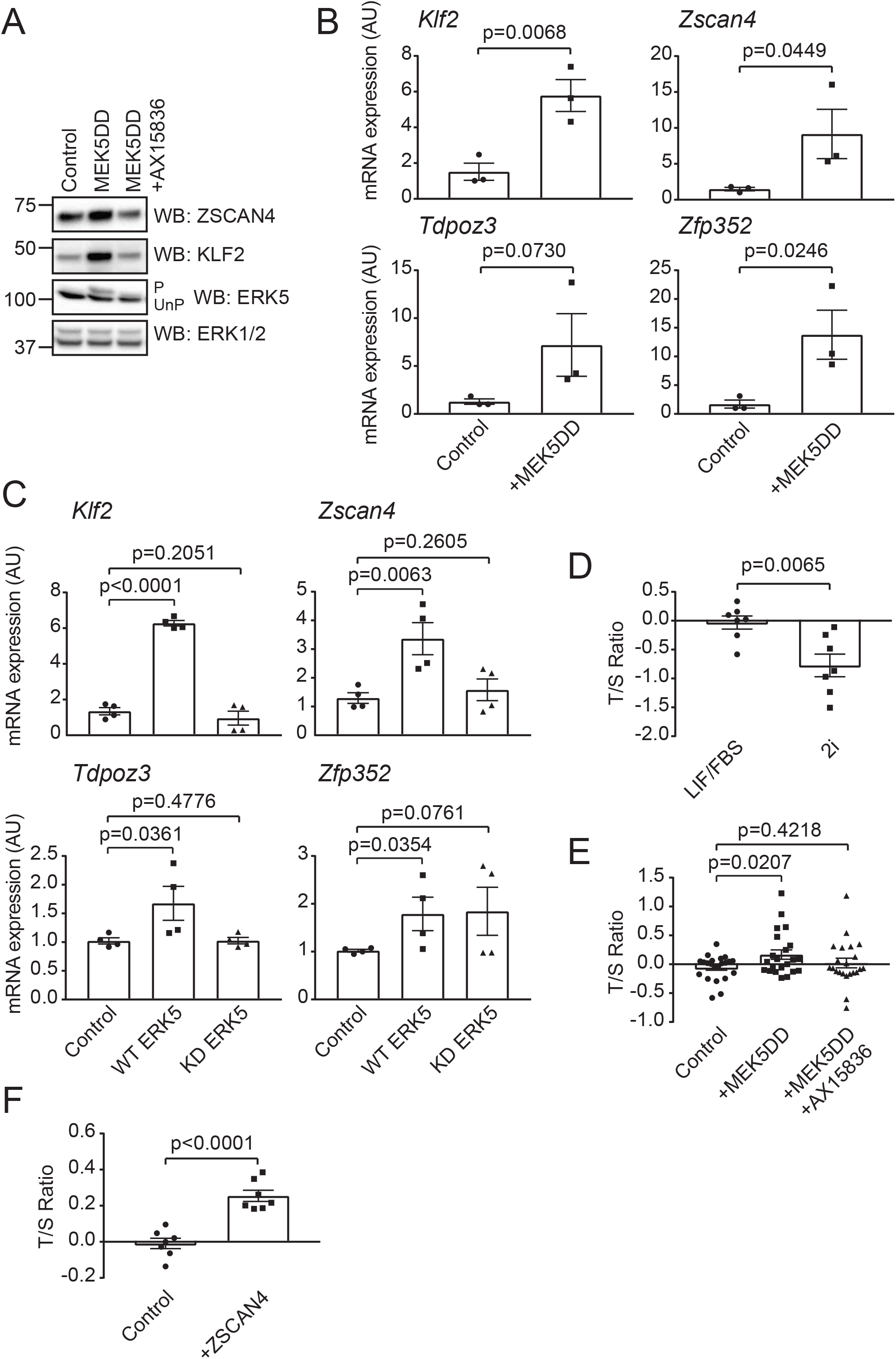
ERK5 promotes ZSCAN4/2-cell stage gene expression and mESC telomere elongation. (A) mESCs were transfected with empty vector control or MEK5DD for (48 h) and treated with either DMSO or 10 μM AX15836 (24 h prior to lysis). ERK5 activation was assessed by band-shift following ERK5 immunoblotting (P = phosphorylated activate ERK5, UnP = unphosphorylated inactive ERK5) and by KLF2 induction. ERK1/2 was used as a loading control. (B) mESCs were transfected with either empty vector or MEK5DD. mRNA levels of *Klf2, Zscan4, Tdpoz3* and *Zfp352* were determined by qRT-PCR. *Klf2* induction is used as a positive control for ERK5 activity. Each data point represents one biological replicate calculated as an average of two technical replicates (n=3). Error bars represent mean ± SEM. Statistical significance was determined by student T test. (C) *Erk5/Mapk7*^-/-^ mESCs were transfected with MEK5DD and either empty vector, wildtype ERK5 or kinase inactive (D200A) ERK5. mRNA levels of *Klf2, Zscan4, Tdpoz3* and *Zfp352* were determined by qRT-PCR. *Klf2* induction is used as a positive control. Each data point represents one biological replicate calculated as an average of two technical replicates (n=3). Error bars represent mean ± SEM. Statistical significance was determined by student T test. (D) Genomic DNA from mESCs maintained in LIF/FBS or 2i media was subjected to qPCR using primers against the telomeric repeats (T) and a single locus control region (S). T/S ratio was calculated to give the average relative telomere length. Each data point represents one biological replicate calculated as the average of three technical replicates (n=7). A power calculation was used to determine sample size, which was randomly selected from biological replicates. Error bars represent mean ± SEM. Statistical significance was determined by student T test. (E) Genomic DNA from mESCs transfected with either empty vector or MEK5DD (48 h) and treated with either DMSO or 10 μM AX15836 (24 h prior to lysis) was collected and subjected to qPCR using primers against the telomeric repeats (T) and a single locus control region (S). T/S ratio was calculated to give the average relative telomere length. Each data point represents one biological replicate calculated as the average of three technical replicates (n=22). A power calculation was used to determine sample size, which was randomly selected from biological replicates. Error bars represent mean ± SEM. Statistical significance was determined by student T test. (F) Genomic DNA from mESCs transfected with either empty vector or ZSCAN4 (48 h) was collected and subjected to qPCR using primers against the telomeric repeats (T) and a single locus control region (S). T/S ratio was calculated to give the average relative telomere length. Each data point represents one biological replicate calculated as the average of three technical replicates (n=7). A power calculation was used to determine sample size, which was randomly selected from biological replicates. Error bars represent mean ± SEM. Statistical significance was determined by student T test.

We then sought to determine the mechanism by which ERK5 signalling drives increased ZSCAN4 protein levels. As ERK5 signalling plays a key role in transcriptional regulation of stem cell-specific genes, we tested whether ERK5 activation induces expression of the *Zscan4* gene cluster. Activation of ERK5 by MEK5DD induces expression of the known target gene *Klf2* (Figure 2B). Similarly, ERK5 activation induces expression of *Zscan4* mRNA (Figure 2B), suggesting that ERK5 promotes transcription of the *Zscan4* gene cluster. Interestingly, ERK5 activation also induces expression of further genes that are specifically expressed at the early embryonic 2C stage *Zfp352* and *Tdpoz3* (Figure 2B), suggesting that ERK5 has a more general function in regulating early embryonic gene expression in mESCs.

Finally, we employed *Erk5/Mapk7*^-/-^ mESCs (Williams et al., 2017) to confirm the role of ERK5 in regulation of *Zscan4*/2C genes. *Erk5*^-/-^ mESCs expressing MEK5DD express a basal level of *Zscan4, Zfp352* and *Tdpoz3* mRNAs (Figure 2C). These levels are strongly increased by expression of wild-type ERK5, in comparison to a kinase inactive mutant (D200A) of ERK5 (ERK5 KD; Figure 2C). Taken together, our data indicate that ERK5 signalling plays a key role in regulating expression of early embryonic genes in mESCs, including the stem cell rejuvenation factor ZSCAN4.

### ERK5-dependent ZSCAN4 induction drives telomere elongation

ZSCAN4 promotes mESC rejuvenation at least in part by promoting telomere maintenance (Amano et al., 2013; Zalzman et al., 2010). Therefore, we asked whether ERK5 signalling to ZSCAN4 impacts on telomere length in mESCs. In order to quantify average telomere length, we employed an assay that determines the ratio of telomeric repeats to non-telomeric DNA (Callicott and Womack, 2006). As proof of principle for this approach, we performed a comparison of telomere length in mESCs cultured in MEK1/2 and GSK3 inhibitors (2i) (Ying et al., 2008) and those cultured in LIF/FCS. mESCs cultured in 2i have been shown to have shortened telomeres compared to LIF/FCS conditions (Guo et al., 2018). Indeed, the telomere:non-telomere ratio is significantly lower for 2i mESCs than LIF/FCS mESCs (Figure 2D), confirming that this assay accurately reports perturbations in telomere length.

We then used this assay to investigate the function of ERK5 signalling in regulating telomere length. ERK5 activation by constitutively-active MEK5DD induces a small but statistically significant increase in telomeric:non-telomeric ratio, which is not observed following co-treatment of mESC with the selective ERK5 inhibitor AX15836 (Figure 2E). Overexpression of ZSCAN4 in mESCs increases the telomeric:non-telomeric ratio (Figure 2F), confirming the key function of ZSCAN4 in regulating telomere length in this context. These data indicate that ERK5 pathway activation drives an increase in telomere length, and this is associated with ZSCAN4 induction.

### The ERK5-KLF2 transcriptional axis controls *Zscan4*/2C gene expression

A key question arising from our results concerns the mechanism by which ERK5 signalling drives ZSCAN4 expression to promote stem cell rejuvenation and telomere maintenance. As shown previously, a major transcriptional target of ERK5 in mESCs is the KLF2 transcription factor (Figure 1F). Therefore, we tested the hypothesis that transcriptional induction of KLF2 is a critical mechanism by which ERK5 signalling drives expression of might be responsible for transcription of *Zscan4* and other 2C genes. To this end, we used CRISPR/Cas9 gene editing to generate mESCs in which expression of wild-type KLF2 is suppressed (*Klf2*^Δ/Δ^ mESCs). In these cells, ZSCAN4 is expressed at a low level, and this is robustly enhanced by expression of wild-type KLF2 (Figure 3A). Expression of *Zscan4, Zfp352* and *Tdpoz3* mRNAs is also increased by re-introduction of wild-type KLF2 (Figure 3B), strongly suggesting that ERK5 signalling regulates expression of ZSCAN4/2C genes via KLF2-dependent transcriptional induction. Consistent with this notion, induction of ZSCAN4 expression by ERK5 activation is blunted in *Klf2*^Δ/Δ^ mESCs when compared to wild-type mESCs expressing endogenous levels of KLF2 (Figure 3C). Taken together, our results indicate that ERK5-KLF2 axis promotes expression of the *Zscan4* and other 2C early embryonic genes.

**Figure 3.**
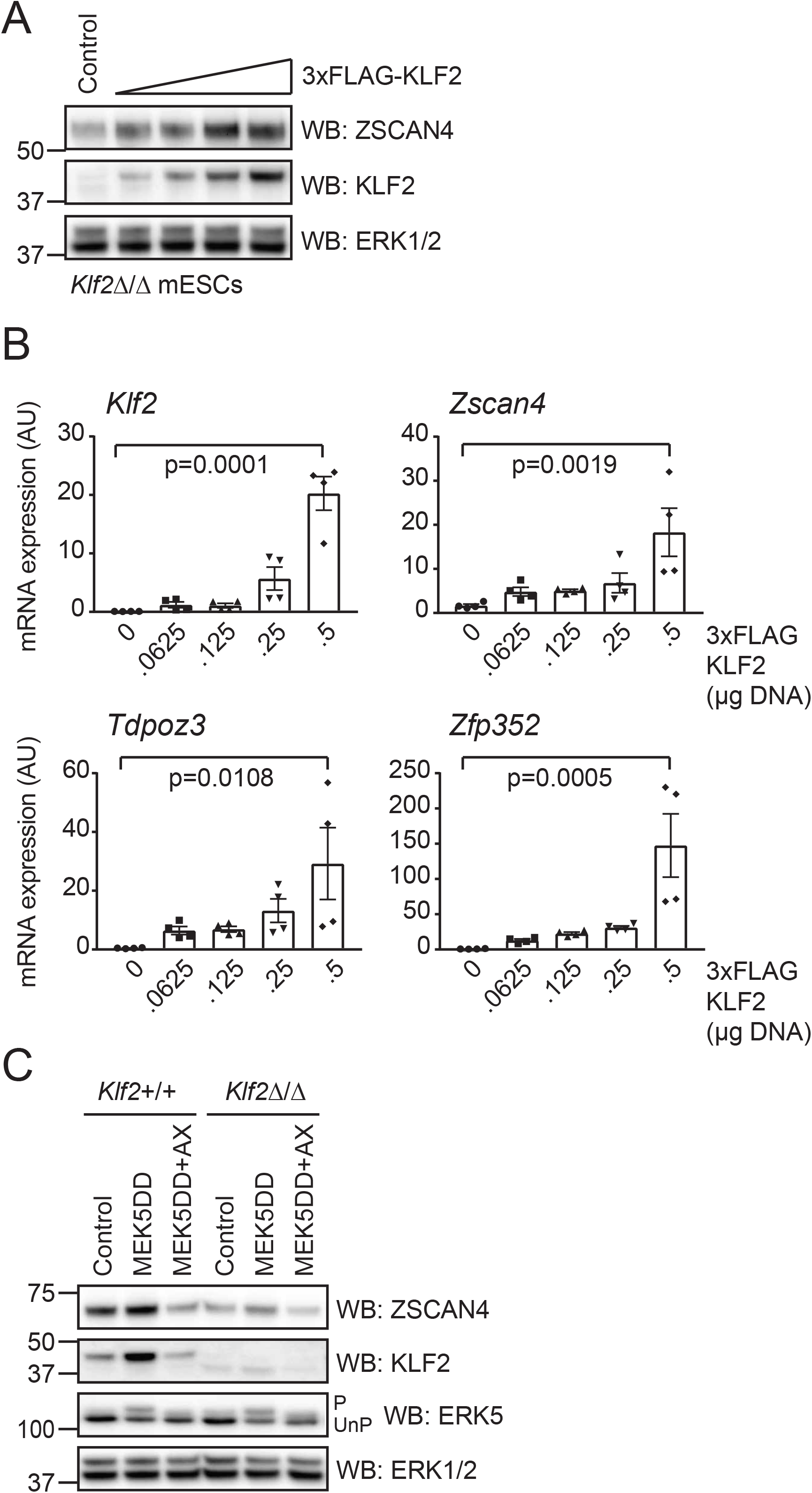
ERK5 signalling to KLF2 regulates ZSCAN4/2-cell stage genes. (A) *Klf2*^Δ/Δ^ mESCs were transfected with either empty vector or increasing concentrations of 3xFLAG-KLF2. Expression levels of KLF2 and ZSCAN4 were determined by immunoblotting. ERK1/2 was used as a loading control. (B) *Klf2*^Δ/Δ^ mESCs were transfected with either empty vector or increasing concentrations of 3xFLAG-KLF2. Levels of *Klf2, Zscan4, Tdpoz3* and *Zfp352* mRNAs were determined by qRT-PCR. Each data point represents one biological replicate calculated as an average of two technical replicates (n=4). Error bars represent mean ± SEM. Statistical significance was determined by one-way ANOVA. (C) Wildtype (*Klf2*^+/+^) and *Klf2*^Δ/Δ^ mESCs were transfected with either empty vector or MEK5DD (48 h) and treated with DMSO or 10 μM AX15836 (24 h prior to lysis). ERK5 activation was assessed by band-shift following ERK5 immunoblotting (P = phosphorylated active ERK5, UnP = unphosphorylated inactive ERK5), and by KLF2 induction. ERK1/2 was used as a loading control.

### ERK5 directly phosphorylates KLF2 at multiple pSer/Thr-Pro motifs

We have demonstrated that KLF2 is a transcriptional target of the ERK5 pathway in regulation of ZSCAN4/2C genes. However, we next explored whether ERK5 might regulate KLF2 via other mechanisms. KLF2 comprises a predicted N-terminal MAP kinase docking motif and phosphomotifs that can be phosphorylated by CMGC family kinases, particularly the MAP kinase family (Figure 4A). This raised the intriguing possibility that ERK5 may additionally regulate KLF2 by direct phosphorylation. We therefore tested whether recombinant ERK5 phosphorylates recombinant KLF2 *in vitro*. ERK5 which is phosphorylated and activated by MEK5 in insect cells and then purified, directly phosphorylates recombinant GST-KLF2 in vitro (Figure 4B). GST-KLF2 phosphorylation is ablated by pre-treatment of the kinase assay with the selective ERK5 inhibitor XMD8-92 (Figure 4B), indicating that trace contaminant kinases are not responsible for KLF2 phosphorylation. Activated ERK5 does not efficiently phosphorylate GST-ATF2 19-96, confirming that the observed phosphorylation is specific for KLF2 and not the GST tag (Figure 4C). Furthermore, we show that ERK5 phosphorylates KLF2 to high stoichiometry of up to 0.5 pmol phosphate/pmol protein (Figure 4D), as might be expected for a *bona fide* ERK5 substrate.

**Figure 4.**
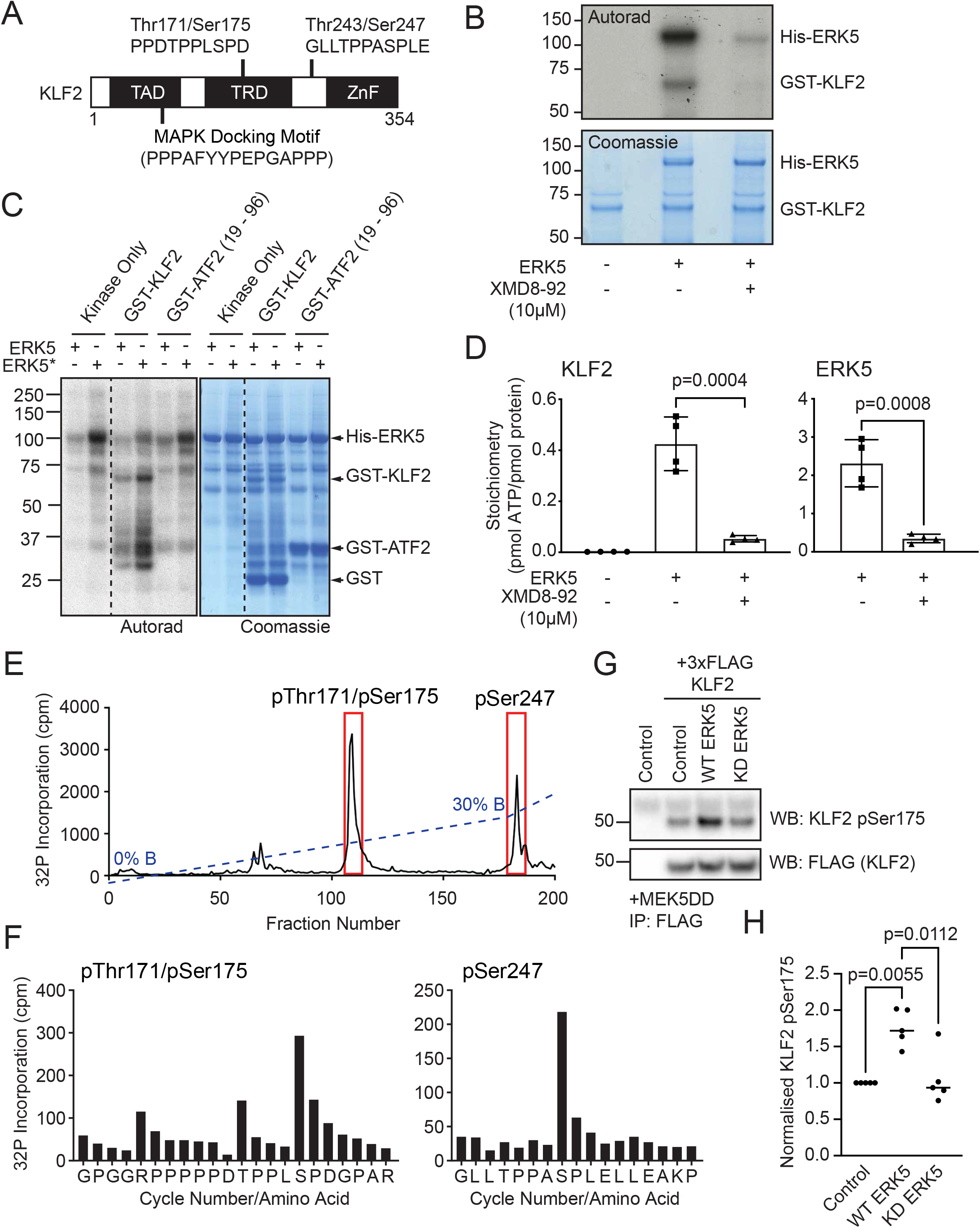
ERK5 directly phosphorylates KLF2 at multiple S/T-P motifs. (A) KLF2 contains a predicted MAP kinase docking motif within its transcriptional activation domain (TAD), and two TPPXSP MAP kinase consensus phosphorylation motifs, one within its transcriptional repression domain (TRD), and one between the TRD and zinc finger (ZnF) domain. (B) Recombinant GST-KLF2 was incubated with [γ-^32^P]-ATP and active recombinant His-ERK5 in the presence or absence of 10 μM of the selective ERK5 inhibitor XMD8-92. KLF2 and ERK5 phosphorylation was assessed by autoradiography and protein loading determined by coomassie blue staining. (C) Recombinant GST-KLF2 or GST-ATF2 (19-96) were incubated with [γ-^32^P]-ATP and either unphosphorylated (ERK5) or phosphorylated active (ERK5*) recombinant His-ERK5. KLF2, ATF2 and ERK5 phosphorylation was assessed by autoradiography and protein loading determined by coomassie blue staining. (D) Recombinant GST-KLF2 was incubated with [γ-^32^P]-ATP and active recombinant His-ERK5 in the presence or absence of 10 μM of the selective ERK5 inhibitor XMD8-92. KLF2 and ERK5 bands were excised from SDS-PAGE and stoichiometry of phosphorylation determined (n=4). Data are represented as mean ± SEM. Statistical significance was determined by student T test. (E) Active recombinant ERK5 was incubated with recombinant KLF2 and [γ-^32^P]-ATP and KLF2 trypsin digested before HPLC analysis. ^32^P phosphorylated KLF2 peptides were identified and the sequence determined by mass spectrometry. (F) Active recombinant ERK5 was incubated with recombinant KLF2 and [γ-^32^P]-ATP. ^32^P phosphorylated tryptic peptides were analysed by mass-spectrometry and Edman sequencing to identify phosphorylated residues. (G) mESCs were transfected with MEK5DD, 3xFLAG-KLF2, and either empty vector, wildtype ERK5 or kinase inactive (D200A) ERK5 (KD). FLAG-KLF2 was isolated by FLAG IP, and FLAG-KLF2 Ser175 phosphorylation and expression assessed by immunoblotting.

We next sought to identify the KLF2 sites which are phosphorylated by ERK5 *in vitro*. To this end, we subjected ERK5 phosphorylated KLF2 to tryptic digestion and HPLC analysis. Radioactive tracing identifies two major peaks of KLF2 phosphorylation (Figure 4E). Edman sequencing shows that the first peak of ERK5 phosphorylation corresponds to a dual phosphorylated Thr171/Ser175 peptide found within the KLF2 transcriptional repression domain, which contains the two Ser/Thr-Pro motifs characteristic of CMGC kinase substrates (Figure 4A). The second peak is a singly phosphorylated Thr243/Ser247 peptide consisting of two further Ser/Thr-Pro motifs towards the KLF2 DNA binding domain (Figure 4F). Interestingly, ERK5 primarily phosphorylates Ser247 but displays little activity towards Thr243 (Figure 3D), in contrast to phosphorylation of the Thr171/Ser175 motif. Therefore, ERK5 directly phosphorylates KLF2 on multiple, distinct Ser/Thr-Pro motifs characteristic of MAP kinase substrates.

We then tested whether ERK5 activity promotes KLF2 phosphorylation at these motifs in mESCs. Using an ectopically-expressed 3xFLAG-tagged KLF2 reporter, which is not subject to transcriptional regulation by ERK5, we monitored KLF2 phosphorylation using a KLF2 Ser175 phospho-specific antibody, which is a major site of ERK5 phosphorylation (Figure 4F). Using this system, we observe that KLF2 is basally phosphorylated at Ser175 (Figure 4G), consistent with KLF2 phosphorylation by other kinases such as ERK1/2 (Yeo et al., 2014). However, co-expression of WT-ERK5, but not a kinase inactive mutant (D200A) of ERK5 (ERK5 KD) drives an increase in KLF2 Ser175 phosphorylation (Figure 4G). These data suggest that ERK5 can also phosphorylate KLF2 at Ser/Thr-Pro motifs in mESCs.

### KLF2 phosphorylation drives Cullin E3 ligase recruitment and ubiquitin-dependent degradation

Our findings then raised the question of mechanism(s) by which ERK5 phosphorylation impacts on KLF2 function. In order to address this, we performed affinity purification mass spectrometry to identify proteins that specifically interact with KLF2 in a manner that is dependent on Ser/Thr-Pro motif phosphorylation. Within the cohort of proteins that are enriched in immunoprecipitates of FLAG-tagged wild-type KLF2 compared to KLF2 with the dual Ser/Thr-Pro phosphorylation motifs mutated (KLF2-S4A) are FBW7, an F-box WD40 containing Cullin substrate adaptor and CUL1, a Cullin family RING-finger E3 ubiquitin ligase (CRL; Figure 5A). Previous studies implicate ubiquitylation in KLF2 regulation in mESCs (Yeo et al., 2014), and FBW7 has been shown to mediate KLF2 ubiquitylation in endothelial cells (Wang et al., 2013; Zhao and Sun, 2013). We validate the interaction between HA-tagged FBW7 and FLAG-KLF2 (Figure 5B) and confirm that mutation of either the Thr171/Ser175 or Thr243/Ser247 phosphorylation motifs (KLF2 2A-N or 2A-C respectively) or all four Ser/Thr-Pro sites (KLF2-4A) disrupts this interaction. This suggests that multiple KLF2 phosphorylation motifs are required for recruitment of FBW7, which is characteristic of phosphorylation dependent ubiquitylation by FBW7 CRL complexes (Koepp et al., 2001). These data therefore indicate that KLF2 phosphorylation promotes recruitment of a CRL E3 ubiquitin ligase. Interestingly, KLF2 Ser/Thr-Pro site mutants appear to show increased expression in mESCs (Figure 5B), suggesting that they are less susceptible to proteasomal degradation.

**Figure 5.**
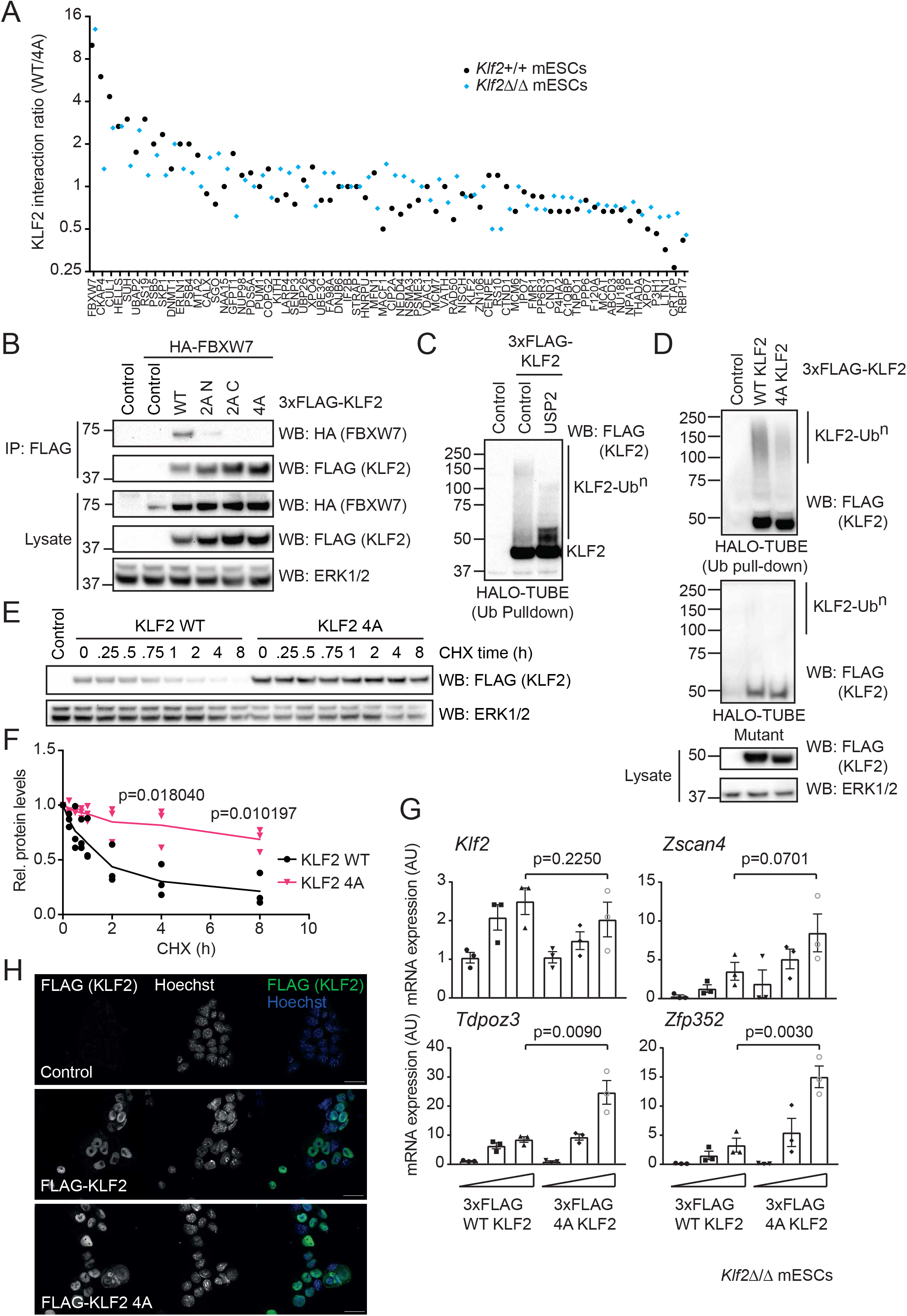
KLF2 phosphorylation drives ubiquitylation and negative feedback control of ZSCAN4/2C genes. (A) Wildtype (*Klf2*^+/+^) or *Klf2*^Δ/Δ^ mESCs were transfected with either wildtype (WT) or T171A/S175A/T243A/S247A mutant (4A) 3xFLAG-KLF2. FLAG-KLF2 was isolated by FLAG IP, and interacting proteins identified by mass spectrometry. Spectral counting was used to rank proteins that preferentially interact with WT KLF2 (interaction ratio >1) or 4A KLF2 (interaction ratio <1). (B) mESCs were transfected with HA-FBXW7 and either empty vector or the indicated 3xFLAG-KLF2 construct. WT = wildtype, 2A N = T171A/S175A, 2A C = T243A/S247A, 4A = T171A/S175A/T243A/S247A. FLAG FLAG-KLF2 was isolated by FLAG IP, and HA-FBXW7 interaction determined by immunoblotting. ERK1/2 was used as a loading control. (C) mESCs were transfected with either empty vector or 3xFLAG-KLF2 and extracts subjected to HALO-TUBE ubiquitin pull-down and incubated in the presence or absence of USP2 deubiquitinase. (D) mESCs were transfected with either empty vector, or 3xFLAG-KLF2 WT or 4A and cell extracts subjected to HALO-TUBE ubiquitin pull-down. Mutant HALO-TUBE beads unable to bind ubiquitin were used to confirm the observed signal can be attributed to ubiquitylated KLF2. ERK1/2 was used as a loading control. (E) mESCs were transfected with either empty vector, or 3xFLAG-KLF2 WT or 4A and treated with cyclohexamide for the indicated time prior to lysis. 3xFLAG KLF2 levels were assessed by immunoblotting, and ERK1/2 levels used as a loading control. (F) Quantification of KLF2 WT and 4A half-life from experimental replicates described in (E). (n=3) Data are represented as mean ± SEM. Statistical significance was determined by student T test. (G) *Klf2*^Δ/Δ^ mESCs were transfected with either empty vector or increasing concentrations of 3xFLAG-KLF2 WT or 4A (48 h). *Klf2, Zscan4, Tdpoz3* and *Zfp352* mRNA levels were determined by qRT-PCR. Each data point represents one biological replicate calculated as an average of two technical replicates (n=3). Data are represented as mean ± SEM. Statistical significance was determined by student T test. (H) mESCs were transfected with either empty vector, or 3xFLAG-KLF2 WT or 4A (24 h), fixed and stained with anti-FLAG antibody to determine KLF2 localisation and Hoechst to stain nuclei. Scale bar = 25 μm.

We then tested the role of dual Ser/Thr-Pro motifs in driving KLF2 ubiquitylation using UBQLN2 Tandem Ubiquitin Binding Element (TUBE) resin to specifically enrich ubiquitylated proteins. FLAG-KLF2 is poly-ubiquitylated in mESCs, as demonstrated by the high-molecular weight species detected in HALO-TUBE pull-downs (KLF2-Ub^n^; Figure 5C). Treatment with the broad-specificity deubiquitinase USP2 abolishes the KLF2-Ub^n^ signal (Figure 5C), confirming that these high-molecular weight species are poly-ubiquitylated KLF2. Furthermore, mutational inactivation of dual Ser/Thr-Pro phosphorylation motifs (KLF2-4A) abolishes KLF2 ubiquitylation (Figure 5D), suggesting that ERK5 phosphorylation of KLF2 at Ser/Thr-Pro motifs promotes KLF2 ubiquitylation. KLF2 phosphorylation and ubiquitylation impacts on stability, as a cycloheximide time-course indicates that wild-type KLF2 is a relatively short-lived protein compared to KLF2 in which the Ser/Thr-Pro motifs are mutated (KLF2-4A; Figure 5E, F). These data therefore implicate ERK5 not only in transcriptional induction of the *Klf2* gene, but also in phosphorylation and resulting proteasomal degradation of the KLF2 protein product.

### ERK5 phosphorylation of KLF2 confers negative feedback regulation of ZSCAN4/2C gene expression

KLF2 transcriptional induction by ERK5 signalling promotes *Zscan4*/2C gene expression. However, ERK5 also directly phosphorylates KLF2 to drive ubiquitylation and degradation. This prompts the hypothesis that KLF2 phosphorylation by ERK5 confers a negative feedback loop to temper *Zscan4*/2C gene expression. In order to test this notion, we investigated the impact of the dual Ser/Thr-Pro phosphorylation motifs on KLF2-dependent transcriptional induction of *Zscan4*/2C genes. As shown previously, expression of wild-type KLF2 drives expression of *Zscan4, Zfp352* and *Tdpoz3* mRNAs (Figure 5G). However, mutation of the dual Ser/Thr-Pro motifs (KLF2-4A) further augments *Zscan4, Zfp352* and *Tdpoz3* expression, suggesting that KLF2 phosphorylation and ubiquitin-mediated turnover functions to suppress KLF2 transcriptional induction of *Zscan4* and other early embryonic genes involved in stem cell rejuvenation. Wild-type KLF2 and KLF2-4A are predominantly, but not exclusively, nuclear localised in mESCs (Figure 5H), indicating that KLF2 phosphorylation does not impact on transcriptional activity by altering sub-cellular localisation. These results therefore suggest that ERK5 signalling promotes ZSCAN4 expression by KLF2 transcriptional induction, and KLF2 function is then tempered by a negative feedback loop dependent upon direct ERK5 phosphorylation and resulting ubiquitylation (Figure 6).

**Figure 6.**
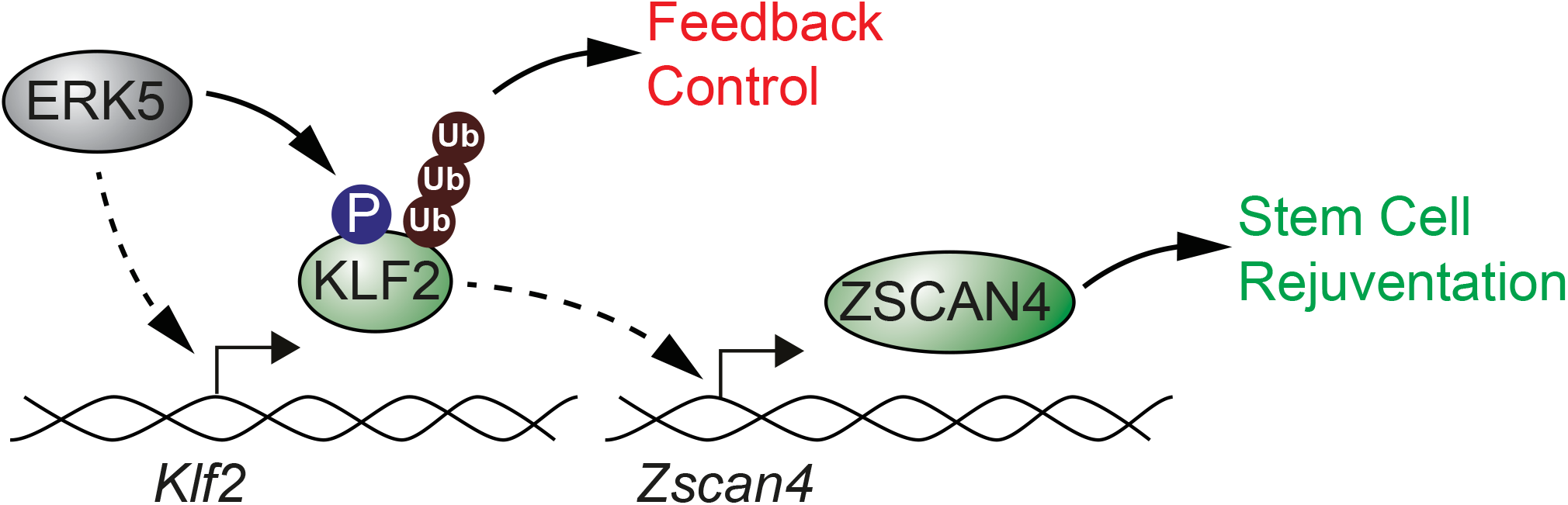
ERK5 signalling to KLF2 confers dynamic regulation of rejuvenation/2-cell stage genes. The ERK5 pathway is subject to complex regulation that we propose plays a critical role in establishing the regulatory dynamics of ZSCAN4 expression. ERK5 activation drives *Klf2* transcription, which drives ZSCAN4/2C gene expression. However, ERK5 also directly phosphorylates KLF2, which negatively regulates KLF2 function by promoting ubiquitylation and degradation. Thus, ERK5 both promotes KLF2 transcriptional induction and negatively regulates KLF2 via direct phosphorylation, generating a dynamic system of amplification followed by negative feedback that accounts for oscillatory ZSCAN4 expression observed in mESCs.

## Discussion

ERK5 is a unique kinase encoding both a kinase domain and a transcriptional activation domain. It was identified as a transcriptional activator of key pluripotency genes in mESCs, including the *Klf2* transcription factor (Williams et al., 2017). However, other functions of ERK5 in regulating gene expression and proteome dynamics in mESCs have not been systematically explored. We use quantitative proteomics to identify the ERK5-dependent proteome, which unveils a function for ERK5 in driving expression of the ZSCAN4 rejuvenation factor and other early embryonic 2-cell stage genes. We show that the mechanism by which ERK5 induces expression of these genes is via KLF2, which is a transcriptional target of the ERK5 signalling pathway.

The ERK5 pathway is subject to complex regulation that we hypothesise play a critical role in establishing the regulatory dynamics of ZSCAN4 expression. Previous work has shown that ERK5 is phosphorylated and activated by MEK5, resulting in ERK5 nuclear translocation (Buschbeck and Ullrich, 2005; Gomez et al., 2016). Once localised to the nucleus, ERK5 induces KLF2 transcription via a dual mechanism involving members of the MEF2 family of transcription factors. This involves direct phosphorylation and activation of MEF2 (Kato et al., 1997) and transcriptional co-activation of MEF2 target genes via the ERK5 C-terminal transcriptional activation domain (Kasler et al., 2000; Sohn et al., 2005). Curiously, we report here that ERK5 also directly phosphorylates KLF2, which in contrast to ERK5 phosphorylation of MEF2 negatively regulates KLF2 function by promoting ubiquitylation and degradation. Thus, ERK5 activation and nuclear translocation presumably enables both ERK5 dependent transcriptional induction of KLF2 and direct phosphorylation and negative regulation of largely nuclear KLF2. We propose this generates a dynamic system of amplification followed by negative feedback, which could account for oscillatory ZSCAN4 expression observed in mESCs (Zalzman et al., 2010).

We have shown that ERK5 signalling to ZSCAN4 promotes telomere maintenance i.e. increased telomere length in mESCs. However, the mechanism by which ZSCAN4 drives telomere maintenance remains a key question. The ZSCAN4 network leads to global demethylation of the mESC genome (Eckersley-Maslin et al., 2016), and reportedly mediates telomere maintenance by suppressing DNA methylation to enable telomere extension via homologous recombination (Dan et al., 2017). This is driven by ubiquitylation and proteasomal degradation of the key maintenance DNA methyltransferase, DNMT1, in a mechanism dependent on the E3 ubiquitin ligase UHRF1 (Dan et al., 2017). Therefore, it will be important to investigate the function of ERK5 signalling to ZSCAN4 in suppression of DNMT1 expression, which could underpin increased telomere elongation activity observed upon ERK5 pathway activation. Of interest, we have shown previously and in this paper that ERK5 signalling acts to suppress expression of *de novo* DNA methyltransferases DNMT3A and DNMT3B (Williams et al., 2016).

Finally, the mechanism by which KLF2 controls ZSCAN4 has not yet been established. KLF2 genome occupancy studies have not identified KLF2 binding sites in the ZSCAN4 gene cluster, although this genomic region is underrepresented in such datasets. Recent data shows that Kruppel-like factors (KZFP factors) related to KLF2 promote activation of transposable element-based enhancers during zygotic genome activation, which in turn induces expression of early embryonic genes during development (Pontis et al., 2019). An exciting possibility is that this transponable element-based mechanism may be responsible for expression of early embryonic genes downstream of the ERK5-KLF2 signalling axis in mESCs.

### Experimental procedures

Many reagents developed for this study are available by request at the MRC-PPU reagents &services website (https://mrcppureagents.dundee.ac.uk/).

### Antibodies and Chemicals

Antibodies used were ERK5 (MRC-PPU Reagents &Services, Dundee), phospho-p38 (Thr180/Tyr182, Cell Signalling), phospho-ERK1/2 (Thr202/Tyr204, Cell Signalling), ERK1/2 (Santa Cruz Biotechnology), GFP (Abcam), KLF2 (Millipore), FLAG (Sigma), ZSCAN4 (Millipore), phospho-KLF2 (DSTT, Dundee), HA (Sigma). Inhibitors used were AX15836 (Tocris), XMD8-92 (N. Gray, Harvard), cycloheximide (Sigma).

### mESC culture, transfection and lysis

CCE mESCs were cultured on gelatin coated plates in media containing LIF, 10% fetal calf serum (Gibco), and 5% knockout serum replacement (Invitrogen) unless otherwise stated. mESCs were transfected with pCAGGS expression vectors using Lipofectamine LTX (Life Technologies), selected with puromycin after 24 h and cultured for the stated times. For CRISPR/Cas9, mESCs were transfected with pX335 and pKN7 (Addgene) and selected, then either lysed or clones isolated. Cell extracts were made in lysis buffer (20 mM Tris [pH 7.4], 150 mM NaCl, 1 mM EDTA, 1% NP-40 [v/v], 0.5% sodium deoxycholate [w/v], 10 mM β-glycerophosphate, 10 mM sodium pyrophosphate, 1 mM NaF, 2 mM Na_3_VO_4_, and Roche Complete Protease Inhibitor Cocktail Tablets).

### Quantitative proteomics

### Protein extraction and digestion

mESCs were washed in PBS and then lysed in 8.5 M urea, 50 mM ammonium bicarbonate (pH 8.0) supplemented with protease inhibitors. Lysate was sonicated using Biosonicator operated at 50% power for 30 s on/off each on ice water bath for 5 min. The lysates were then centrifuged at 14,000 rpm for 10 mins at 4 °C and supernatants collected. Protein concentration of the lysate was determined by BCA protein assay. Proteins were reduced with 5 mM DTT at 55 °C for 30 min and cooled to room temperature. Reduced lysates were then alkylated with 10 mM iodoacetamide at room temperature for 30 mins in the dark. The alkylation reaction was quenched by the addition of another 5 mM DTT. After 20 min of incubation at room temperature, the lysate was digested using Lys-C with the weight ratio of 1:200 (Enzyme/lysate) at 37 °C for 4 h. The samples were further diluted to 1.5 M Urea with 50 mM ammonium bicarbonate (pH 8.0), and the sequencing-grade trypsin was added with the weight ratio of 1:50 (enzyme/lysate) and incubated overnight at 37 °C. The digest was acidified to pH 3.0 by addition of TFA to 0.2% and gently mix at room temperature for 15 min; the resulting precipitates were removed by centrifugation at 7100 RCF for 15 min. The acidified lysate was then desalted using a C18 SPE cartridge (Waters) and the eluate was aliquoted into 100 μg and dried by vacuum centrifugation. To check the digests, 1 µg of each sample was analysed by mass spectrometry prior to TMT labelling.

### TMT labelling and high pH reverse phase fractionation

100 µg of peptide from each sample was re-suspended into 100 mM Triethylammonium bicarbonate buffer (pH 8.5). Then 0.8 mg of TMT tag (Thermo) dissolved in 41 µl of anhydrous acetonitrile was transferred to the peptide sample and incubated with 60 min at room temperature. The TMT labelling reaction was quenched with 5% hydroxylamine. 1 µg of each labelled sample was analysed by mass spectrometry to assess the labelling efficiency before pooling. After checking the labelling efficiency, the TMT-labelled peptides were mixed together and dried by vacuum centrifugation. After dryness, the mixture of TMT-labelled peptides was dissolved into 0.2% TFA and then desalted using a C18 SPE cartridge. The desalted peptides were subjected to orthogonal basic pH reverse phase fractionation, collected in 96-well plate and consolidated for a total of 20 fractions for vacuum dryness.

### LC-MS/MS analysis

Each fraction was dissolved in 0.1% FA and quantified by Nanodrop. 1 µg of peptide was loaded on C18 trap column at a flow rate of 5 μl/min. Peptide separations were performed over EASY-Spray column (C18, 2 µm, 75 mm × 50 cm) with an integrated nano electrospray emitter at a flow rate of 300 nl/min. The LC separations were performed with a Thermo Dionex Ultimate 3000 RSLC Nano liquid chromatography instrument. Peptides were separated with a 180 min segmented gradient as follows: 7%∼25% buffer B (80% ACN/0.1% FA) in 125 min, 25%∼35% buffer B for 30 min, 35%∼99% buffer B for 5 min, followed by a 5 min 99% wash and 15 min equilibration with buffer A (0.1% FA).

Data acquisition on the Orbitrap Fusion Tribrid platform with instrument control software version 3.0 was carried out using a data-dependent method with multinotch synchronous precursor selection MS3 scanning for TMT-9plex tags. The mass spectrometer was operated in data-dependent most intense precursors Top Speed mode with 3 s per cycle. The survey scan was acquired from m/z 375 to 1500 with a resolution of 120,000 resolving power with AGC target 400,000. The maximum injection time for full scan was set to 60 ms. For the MS/MS analysis, monoisotopic precursor selection was set to peptide. AGC target was set to 50,000 with the maximum injection time 120 msec. Charge states unknown and 1 or higher than 7 were excluded. The MS/MS analyses were performed by 1.2 m/z isolation with the quadrupole, normalised HCD collision energy of 37% and analysis of fragment ions in the Orbitrap using 15,000 resolving power with auto normal range scan starting from m/z 110. Dynamic exclusion was set to 60 s. For the MS3 scan, the MS3 precursor population from MS2 scan ranging from m/z 300-100 was isolated using the SPS waveform and then fragmented by HCD. The HCD normalized collision energy was set to 65. The MS3 scan were acquired from m/z 100 to 500 with a resolution of 50,000 and AGC target 50,000/ The maximum injection time for full scan was set to 86 ms.

### Data processing and spectra assignment

Data from the Orbitrap Fusion were processed using Proteome Discoverer Software (version 2.2). MS2 spectra were searched using Mascot against a UniProt Mouse database appended to a list of common contaminants (10,090 total sequences). The searching parameters were specified as trypsin enzyme, two missed cleavages allowed, minimum peptide length of 6, precursor mass tolerance of 20 ppm, and a fragment mass tolerance of 0.05 Daltons. Oxidation of methionine and TMT at lysine and peptide N-termini were set as variable modifications. Carbamidomethylation of cysteine was set as a fixed modification. Peptide spectral match error rates were determined using the target-decoy strategy coupled to Percolator modeling of positive and false matches. Data were filtered at the peptide spectral match-level to control for false discoveries using a q-value cut off of 0.01, as determined by Percolator. For quantification, the signal-to-noise values higher than 10 for unique and razor peptides were summed within each TMT channel, and each channel was normalized with total peptide amount. Quantitation was further performed by adjusting the calculated p-values according to Benjamini-Hochberg. The significance regulated proteins with p-value less than 0.05 were further manually investigated with the standard deviations of biological replicates.

### Immunofluorescence microscopy

mESCs were seeded on gelatin-coated coverslips and transfected as required as described above. After 24 h, mESCs were fixed with 4% PFA (w/v) in PBS, before being permeabilised in 0.5% Triton X-100 in PBS (v/v) for 5 mins at room temperature. Coverslips were then blocked with 1% Fish gelatin in PBS (w/v), and incubated with primary antibodies for 2 h at room temperature in a humid chamber. After three washes with PBS, secondary antibodies conjugated to fluorophores were diluted 1:500 in blocking buffer and incubated on coverslips for 1 h at room temperature in a humid chamber. Where cytoskeleton was being observed, Actin Red 555 reagent was added to the secondary antibody mix. After three washes with PBS, 0.1 μg/ml Hoescht was incubated with the coverslips for 5 mins at room temperature in a humid chamber to stain nuclei. After three more washes with PBS, coverslips were mounted onto cover slides using Fluorsave reagent (Millipore). Images were taken using a Leica SP8 confocal microscope and processed using FIJI and Photoshop CSC software (Adobe).

### RNA extraction and quantitative PCR

RNA was extracted using the OMEGA total RNA kit and reverse transcribed using iScript reverse transcriptase (Bio-Rad). qPCR was performed using TB Green Premix Ex Taq (Takara). The ΔCt method using *Gapdh* as a reference gene was used to analyse relative expression and the 2-ΔΔCt (Livak) method used to normalize to control. Primers used are listed below.

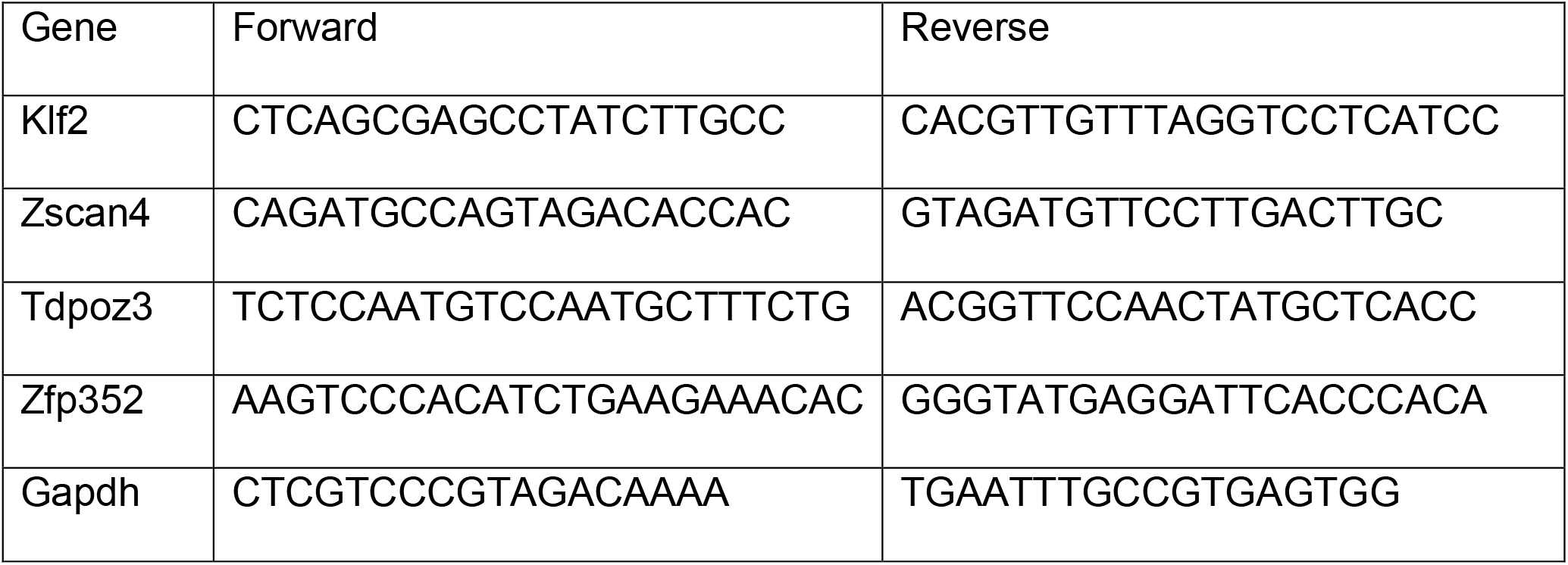

### Recombinant ERK5 kinase assay

200 ng pure active ERK5 was incubated with the indicated inhibitor in 50 mM Tris-HCl (pH 7.5), 0.1 mM EGTA, and 1 mM 2-mercaptoethanol. The reaction was initiated by adding 10 mM magnesium acetate, 100 μM [γ-^32^P]-ATP (500 cpm/pmol), and 5ug of the indicated substrate (GST-KLF2 or GST) and incubated at 30°C for 20 min. The assay was terminated by addition of 4xLDS. Stoichiometry of KLF2 and ERK5 phosphorylation was quantified by excising the specific band from a coomassie blue stained SDS-PAGE gel and ^32^P incorporation measured by Cerenkov counting.

### KLF2 phosphorylation site mapping

GST-KLF2 purified from E. coli was incubated with activated ERK5 for 30 mins at 30 °C in kinase assay reaction buffer with 25000 cpm/pmol ATP in a total reaction volume of 25 μl. The reaction was terminated by boiling with 1x LDS sample buffer plus reducing agent and subjected to electrophoresis on a NuPAGE 4-12% gel. The gel was coomassie stained and the band corresponding to substrate excised and cut into small cubes. From this stage, all steps were conducted in a laminar flow hood to prevent keratin contamination. Gel pieces were washed (10 mins, 500 μl per wash) sequentially on a Vibrax shaking platform with ddH2O, 50% acetonitrile/ddH2O, 0.1 M NH_4_HCO_3_ and 50% acetonitrile/50 mM NH_4_HCO_3_. Gel pieces were reduced further 10 mM DTT at 65 °C for 45 mins and alkylated with 50 mM iodoacetamide (Incubated in the dark for 30 mins at room temperature). Gel pieces were then washed with 50 mM NH_4_HCO_3_ and 50% acetonitrile/50 mM NH_4_HCO_3_. Once colourless, gel pieces were shrunk with acetonitrile for 15 mins, the supernatant aspirated and gel pieces dried using a SpeedVac. Dry gel pieces were swollen with 25 mM Triethylammonium bicarbonate containing 5 μg/ml of Trypsin and incubated at 30 °C overnight on a Thermomixer. After trypsinisation, an equivalent volume of acetonitrile was added to the digested gel pieces and incubated for 15 mins more. Supernatant was transferred to a clean tube and dried in a SpeedVac. Further extraction of peptides from the gel pieces was achieved using 100μl 50% acetonitrile with 2.5% formic acid, and incubated for 15 mins at room temperature. This supernatant was combined with the dry peptides and again dried in a SpeedVac.

Digestion removed >95% of the ^32^P from the gel pieces to the recovered dried peptides. Peptides were fractionated on a HPLC column (Vydac C18) equilibrated in 0.1% (w/v) trifluoroacetic acid (TFA), using a linear acetonitrile gradient at a flow rate of 0.2 ml/min. 100 μl fractions were collected and analysed by LC-MS/MS. The data from this was searched using Mascot (matrixscience.com) to identify Phospho (Ser/Thr), Phospho (Tyr), Oxidation (Met) and Dioxidation (Met). Radioactive fractions were sent to solid-phase Edman degradation for phosphorylation site identification using an Applied Biosystems 494C sequencer of the peptide coupled to Sequelon-AA membranes (Applied Biosystems).

### HALO-TUBE ubiquitin pulldown assay

HALO-tagged UBQLN1 Tandem Ubiquitin Binding Element (TUBE) (MRC-PPU R&3×0026;S DU23799) is a tetramer of the UBQLN1 UBA ubiquitin binding domain (aa536-589) provided by Dr. A. Knebel (MRC-PPU, Dundee). A mutant TUBE incapable of binding ubiquitin (M557K, L584K) was also used. HALO-TUBE beads were prepared by washing 1 ml packed HALO resin three times with HALO wash buffer (50 mM Tris/HCl (pH7.5), 0.5 M NaCl, 1% Triton-X100 (v/v)), resulting in a 1:4 slurry. The slurry was combined with 7 mg of HALO-TUBE or HALO-mutant TUBE and incubated on a rotating wheel at 4 °C overnight. After five washes in 10 ml HALO wash buffer and then in 10 ml HALO storage buffer (50 mM Tris/HCl (pH7.5), 150 mM NaCl, 0.1 mM EGTA, 270 mM Sucrose, 0.07% β-mercaptoethanol (v/v)), the beads were stored as a 20% slurry at 4 °C and washed three times in lysis buffer immediately prior to use. To analyse ubiquitylation of a protein of interest, 10 μl washed packed beads were incubated with 1 mg clarified cell lysate for 3 h at 4 °C with shaking. After three washes with lysis buffer, beads were analysed by SDS-PAGE and immunoblotting.

### Telomere qPCR assay

Average telomere length was determined using qPCR and quantifying the ratio of telomeric DNA repeats to a single-copy gene (acidic ribosomal phosphoprotein PO (36B4) gene. Cells were seeded in 12-well plates, genomic DNA extracted using DNeasy Blood and Tissue Kit (QIAGEN) according to the manufacturer’s instructions. Primers were supplied by Sigma-Aldrich and noted below.

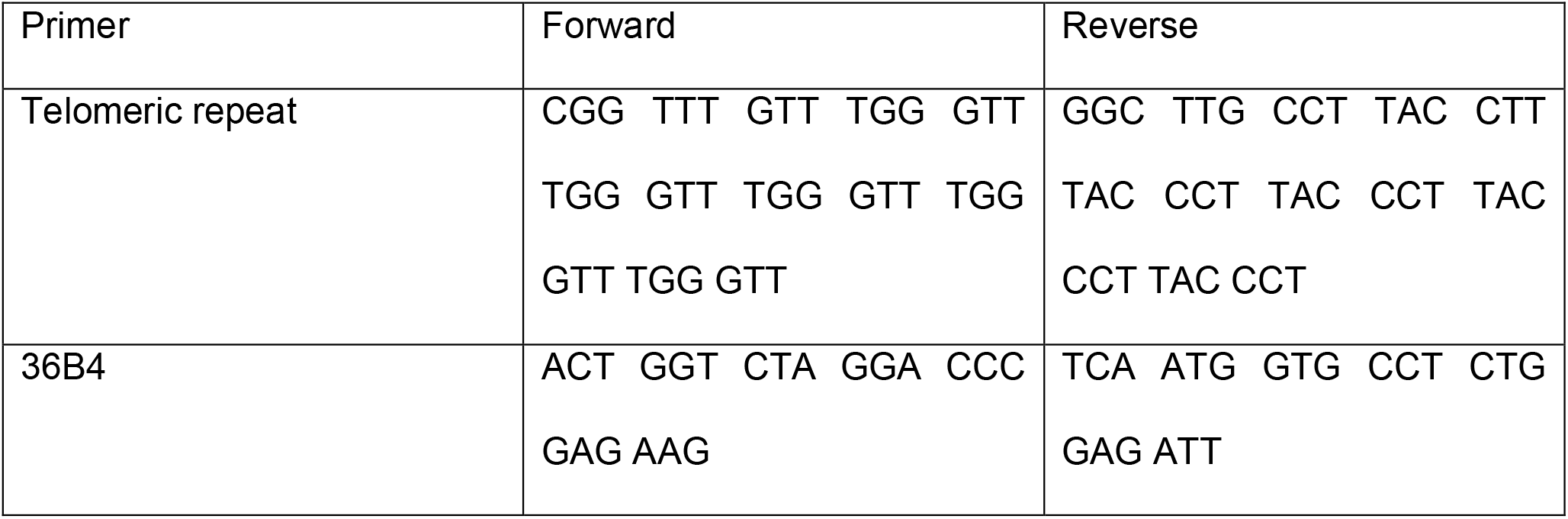

Each reaction for the telomere portion of the assay included 12.5 μl SYBR Green PCR Master Mix (Takara), 300 nM each of the forward and reverse primers, 20 ng genomic DNA, and enough double-distilled H_2_O to yield a 25-μl reaction. Three 20 ng samples of each DNA were placed in adjacent wells of a 96-well plate. An automated thermocycler (Prism 7000 Sequence Detection System, Applied Biosystems) was used with the following reaction conditions: 95 °C for 10 min followed by 30 cycles of data collection at 95 °C for 15 s and a 56 °C anneal–extend step for 1 min. Each reaction for the 36B4 portion contained 12.5 μl SYBR Green PCR Master Mix (Takara), 300 nM forward primer, 500 nM reverse primer, 20 ng genomic DNA and enough double-distilled H_2_O to yield a 25-μl reaction. Three 20 ng samples of each DNA were placed in adjacent wells of a 96-well plate. The thermocycler reaction conditions were: 95 °C for 10 min, followed by 35 cycles of data collection at 95 °C for 15 s, with 52 °C annealing for 20 s, followed by extension at 72 °C for 30 s. Telomere and 36B4 reactions were performed on separate plates so to minimise variation due to location in plate, each sample was loaded into corresponding positions on each plate, such that the two plates had the same layout. Analysis was performed by subtracting 36B4 Ct value from corresponding Telomere Ct value and normalising to the average of the control conditions. These log ratios were then plotted using GraphPad Prism software, which was used to perform statistical tests. Original protocol for telomere qPCR protocol and data analysis is reported in (Callicott and Womack, 2006).

### Statistical analysis

Data are presented as the average with error bars indicating standard error of the mean (SEM). Statistical significance of differences between experimental groups was assessed using a Student’s t test. Differences in averages were considered significant if p < 0.05. Representative western blots are shown. Power calculations for telomere qPCR assays were performed using an online tool (http://powerandsamplesize.com/Calculators/Compare-2-Means/2-Sample-Equality), with the following parameters: sample size nB = number of observed measurements for control group, Power = 0.80, p value = 0.05, Group A mean = mean of observed measurements for test group, Group B mean = mean of observed measurements for control group, Standard deviation = calculated standard deviation based on observed measurements for test group, Sampling ratio = 1. Total sample size was then read off the graph at Power = 0.80 and observed test group mean. Total sample size was divided by 2 to give number of samples required for each group, and replicates were randomly selected for each sample group. A student T test was performed on these sample groups to determine significance.

## Data availability

The mass spectrometry proteomics data have been deposited to the ProteomeXchange Consortium via the PRIDE partner repository with the dataset identifier PXD024679. Other data generated and/or analysed during the current study are available from the corresponding authors on request.

## Acknowledgements

We thank the MRC-PPU Reagents & Services protein production team for recombinant proteins used in this study. H.A.B., D.R-S. and S.M were supported by a MRC DTP PhD studentship, C.A.C.W. was supported by a MRC-PPU prize studentship, H.Z., R.T. and R.G. were supported by the MRC-PPU core operating grant, R.F-A. was supported by the MRC-PPU and L.M. was supported by the a University of Dundee MSc. G.M.F is supported by a Wellcome Trust/Royal Society Sir Henry Dale Fellowship (211209/Z/18/Z) and a MRC New Investigator Research Grant (MR/N000609/1).

## Author contributions

H.A.B., C.A.C.W., D.R-S., S.M., R.F-A., L.M. and N.D. performed biochemical and cell-based experiments, analysed data and prepared figures. H.Z. performed mass-spectrometry and data analysis, R.T. performed molecular cloning and R.G. performed phosphosite identification. J.P., M.T., J.L. and M.S. provided technical and conceptual advice and expertise. G.M.F. coordinated the study and wrote the paper with input from all authors.

## Competing interests statement

The author(s) declare that there are no competing interests.

## Notes

http://www.ebi.ac.uk/pride/archive/projects/PXD024679

### Competing Interest Statement

The authors have declared no competing interest.

